# Robust blind spectral unmixing for fluorescence microscopy using unsupervised learning

**DOI:** 10.1101/797993

**Authors:** Tristan D. McRae, David Oleksyn, Jim Miller, Yu-Rong Gao

## Abstract

Due to the overlapping emission spectra of fluorophores, fluorescence microscopy images often have bleed-through problems, leading to a false positive detection. This problem is almost unavoidable when the samples are labeled with three or more fluorophores, and the situation is complicated even further when imaged under a multiphoton microscope. Several methods have been developed and commonly used by biologists for fluorescence microscopy spectral unmixing, such as linear unmixing, non-negative matrix factorization, deconvolution, and principal component analysis. However, they either require pre-knowledge of emission spectra or restrict the number of fluorophores to be the same as detection channels, which highly limits the real-world applications of those spectral unmixing methods. In this paper, we developed a robust and flexible spectral unmixing method: Learning Unsupervised Means of Spectra (LUMoS), which uses an unsupervised machine learning clustering method to learn individual fluorophores’ spectral signatures from mixed images, and blindly separate channels without restrictions on the number of fluorophores that can be imaged. This method highly expands the hardware capability of two-photon microscopy to simultaneously image more fluorophores than is possible with instrumentation alone. Experimental and simulated results demonstrated the robustness of LUMoS in multi-channel separations of two-photon microscopy images. We also extended the application of this method to background/autofluorescence removal and colocalization analysis. Lastly, we integrated this tool into ImageJ to offer an easy to use spectral unmixing tool for fluorescence imaging. LUMoS allows us to gain a higher spectral resolution and obtain a cleaner image without the need to upgrade the imaging hardware capabilities.

## Introduction

Two-photon laser scanning microscopy (2PLSM) offers many advantages for imaging cell dynamics in live animals with deeper tissue penetrations, 3D contrast and resolution, and reduced phototoxicity [1,2]. The majority of *in-vivo* 2PLSM studies so far have relied on single or dual color imaging which highly limits the cell populations and physiological components that can be studied at one time [3–5]. To identify and characterize complex biological mechanisms, multiple cell types or intracellular processes need to be visualized simultaneously. Adapting 2PLSM for simultaneous multi-fluorophore detection has presented a challenge due to the widely overlapping two-photon absorption spectra of commonly used fluorescent markers [6–8] as well as the high expense of incorporating multiple two-photon laser lines. Imaging specimens with a greater number of fluorescent labels is usually confronted with the bleed-through or cross-talk of fluorescence emissions. These spectral mixing artifacts often complicate the interpretation of experimental results with ambiguous discriminations, particularly if colocalization of fluorophores is under investigation or quantitative measurements are necessary. Therefore, a reliable and clean separation of different fluorescence labels is required for analysis and quantifications, and a flexible approach to overcome the hardware limitations on the number of fluorophores that can be simultaneously imaged is desired.

There are a wide variety of computational approaches commonly used by biologists for spectral unmixing with their own advantages and limitations. Fluorescence signals were first modeled as a linear combination of measured reference spectra of all involved fluorochromes, and linear unmixing was introduced for spectral unmixing in the fluorescence microscopy domain [9,10]. This algorithm extracts the weight of each individual spectrum with the weight proportional to the fluorophore’s concentration [11,12]. Linear unmixing is advantageous in the way that it is well suited for resolving spectra from pixels that have a mixed contribution from different fluorophores, as it calculates the best linear fit of any combination of fluorescent spectra in an individual pixel. The method has been widely applied in different imaging modalities since then [12–16]. However, the spectra of the contributing fluorophores may change nonuniformly due to the distortion by the complex tissue environment [17], and the assumption of superposition may be inappropriate in the presence of non-linear effects such as quenching, photobleaching, and two-photon absorption. To be solvable, linear unmixing also assumes that the number of detection channels be at least equal to the number of fluorophores which requires more advanced hardware settings such as tunable filters to detect more dyes [10,18], highly limiting the number of different labels that can be unambiguously identified in an image. In addition, the method also requires prior knowledge of the reference spectrum for a given dye, which is instrument specific and hard to measure. Following linear unmixing theory, many other methods have been introduced. Non-negative matrix factorization (NMF) considers the non-negative characteristics of the fluorophore contributions [19–21], which has the advantage that prior knowledge of emission spectra is not needed, and has also been used for autofluorescence and background removal [22–24]. It is limited, however, in that it cannot be applied to situations when fluorophores outnumber detection channels. The other main problem of this approach is that there can be multiple equally valid, yet significantly different solutions. Sometimes prior knowledge about spectra is still needed to reduce the ambiguity [19]. Another unmixing method, spectral deconvolution [25], requires the acquisition of the spectral signature of each fluorophore by manually selecting the region of interest which is laborious, and requires unambiguous and exclusive expression of fluorescent labels. The method will not work when, in addition to bleed-through, there is significant cross-talk between fluorophores. Another recently developed method used for two-photon imaging, similarity unmixing [26], can work for any number of fluorophores but still requires detailed knowledge of fluorophore emission spectra and can fail when actual emissions deviate from their theoretical ideals or there are colocalized fluorophores.

Therefore, to improve the flexibility and applicability of multi-channel fluorescence imaging spectral unmixing, we looked for methods that do not need spectra information and are not restricted by the number of detection channels. Unsupervised learning is a class of machine learning techniques that find patterns directly from unlabeled data [27,28]. By taking advantage of the ability of unsupervised learning algorithms to automatically “learn” to identify features from raw images, we here investigated clustering based unsupervised learning in blindly unmixing channels of multi-color 2PLSM images: Learning Unsupervised Means of Spectra (LUMoS). Similar clustering methods have been applied for spectral unmixing in the remote sensing field [29–31], but never to fluorescence microscopy. By assuming the discrete labeling of biological structures, our model uses *k*-means clustering to “learn” the relationships between pixels from the raw image, and search for their intensity patterns to re-classify each pixel into a unique fluorophore group [32]. We emphasize that LUMoS requires neither the knowledge of emission spectra nor a greater or equal number of detection channels than fluorophores, which highly expands the capability of two-photon imaging. We have successfully demonstrated the ability of LUMoS to cleanly separate out up to 6 fluorophores in biological samples imaged by a 2PLSM system with only 4 detectors. Synthetic results demonstrated the accuracy and power of LUMoS in separating more fluorophores under the challenging conditions of unbalanced structure size and low signal-to-noise ratio (SNR). The method can be easily translated to images acquired by other fluorescence imaging modalities such as confocal to create a clean representation of the fluorophores in the sample for quantitative analysis.

## Materials and methods

### Sample preparations

For N_fluorophores_ = N_detectors_ unmixing studies, FluoCells Prepared Slide #1 (F36924, Invitrogen, Carlsbad, CA) was used. Fixed bovine pulmonary artery endothelial (BPAE) cells were stained with a combination of fluorescent dyes. DAPI was used to label the nuclei, F-actin was stained using Alexa Fluor488 (AF488) phalloidin, and mitochondria were labeled with MitoTracker Red CMXRos.

For N_fluorophores_ > N_detectors_ beads unmixing studies, particles of different sizes and colors were mixed and prepared on a glass slide and covered with a #1 cover slip for imaging. The commercial beads were either surface conjugated with standard fluorophores or polymerized with an organic dye within. The emission (EM) peak was either determined by the online spectra-viewer for a standard fluorophore or provided by the nanoparticle company (Spherotech Inc., Lake Forest, IL) for an organic dye. A combination of 5 different beads was used: Light Yellow (LY, FP-2045-2, Spherotech Inc.): 1.97µm in diameter, 450nm peak EM; FITC (ECFP-F1, Spherotech Inc.): 3.27µm in diameter, 515nm peak EM; PE (ECFP-F2, Spherotech, Inc.): 3.4µm in diameter, 575nm peak EM; Purple (FP-2062-2, Spherotech Inc.): 2.37µm in diameter, 620nm peak EM; APC (345036, BD Biosciences, San Jose, CA): 6µm in diameter, 660nm peak EM.

For N_fluorophores_ > N_detectors_ Colorful Cell unmixing studies, a plasmid encoding 6 independent transcription units driving expression of different fluorescent proteins to distinct intracellular compartments, Colorful Cell [33], was a gift from Pierre Neveu (RRID:Addgene_62449; http://n2t.net/addgene:62449; Addgene, Watertown, MA). The 6 fluorescent proteins were TagBFP trimer fused to a nuclear localization sequence, Cerulean trimer fused to a plasma membrane targeting sequence, AzamiGreen fused to a mitochondrial localization sequence, Citrine fused to a Golgi targeting sequence, mCherry fused to an endoplasmic reticulum retention sequence, and iRFP670 fused to a peroxisome localization sequence. Plasmid DNA was transiently transfected into HEK293T (Pheonix) cells by calcium phosphate precipitation and assayed 48-72 hours later. By flow cytometry 40-70% of cells were expressing the transfected plasmid. For imaging, wet mounts of single cell suspensions containing 100,000 live cells/10 µL were prepared and imaged immediately.

For the colocalization experiments, CD28-deficient, DO11.10 T cells were retrovirally transduced with CD28 fused at the C terminus to YFP or to Cerulean either separately or together. T cells were then mixed with stably transfected antigen-presenting cells (APCs) expressing MHC class II, ICAM-1, and CD80 that were or were not preloaded with 2.0 µg/ml OVA peptide for 1 hour at 37°C, and pelleted at Rcf 2000 for 20 sec. The pellet was incubated at 37° C for 10 min, resuspended and plated on poly-L–lysine coated cover slips for imaging [34].

### Two-photon imaging

All images were collected by an Olympus FVMPE-RS system (Olympus, Center Valley, PA) using Olympus 25× water objective (XLPLN25XWMP2, 1.05NA). The system was equipped with two two-photon lasers: Spectra-Physics InSightX3 (680nm-1300nm, Spectra-Physics, Santa Clara, CA) and Spectra-Physics MaiTai DeepSee Ti:Sapphire laser (690nm-1040nm). There were four Photon Multiplier Tubes (PMTs) and two filter cubes for multi-color imaging. Galvanometer scanners were used for scanning. PMT gains for all imaging were used between 500 and 650 a.u. in the Olympus Fluoview software. The system schematic is shown in S1 Fig (the Blue/Green, and Red/fRed filter cubes setup is shown).

For N_fluorophores_=N_channels_ unmixing studies, FluoCells Prepared Slide #1 was imaged using MaiTai laser at 780nm to excite DAPI, AF488, and MitoTracker Red in the BPAE cells. 3D 512×512 pixel images were collected with 0.5µm per z step. For N_fluorophores_>N_channels_ beads unmixing studies, multi-color beads slide was imaged using InSightX3 laser at 1000nm and MaiTai laser at 800nm simultaneously. 2D 512×512 pixel images were collected. For N_fluorophores_>N_channels_ Colorful Cell separation studies, Colorful Cell slide was imaged using InSightX3 laser at 1050nm and MaiTai laser at 840nm sequentially with 1024×1024 pixels in x-y and 0.5µm per z step. Blue/Green cube (420-460nm/495-540nm) and red/fRed cube (575-630nm/645-685nm) were used for the above imaging. For colocalization studies, Cerulean, YFP, or Cerulean + YFP labeled cell slides were imaged with 800×800 pixels in x-y and 0.5µm per z step using InSightX3 laser at 970nm and MaiTai laser at 860nm sequentially. CFP/YFP cube (420-500nm/519-549nm) was used for this colocalization experiment.

### Data pre-processing

Depending on the content of the input image, it may be appropriate to group together pixels with different net intensities but similar ratios of intensities in different z-planes. This could be necessary in fluorescence microscopy, and especially 2PLSM, in which there usually are signal intensity differences across imaging depths. This can be accounted for by dividing the intensity of a pixel *x* in each channel *c* by the overall sum of that pixel intensities across all the channels:

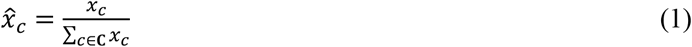

where *x*_*c*_ is the raw intensity of pixel *x* in channel *c*, 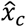 is the scaled intensity of pixel *x* in channel *c* and **C** is the set of all input channels. This step is not always desirable, as in some cases pixels with the same intensity ratios but different raw intensities may actually represent different structures.

To prevent the clustering algorithm from being biased by signal intensity differences and variations between channels, the brightness and contrast of input data were normalized to be relatively spherical distributions before clustering. Normalization also makes *k*-means initialize with better centroid choices and run faster with fewer iterations to converge [32,35]. Therefore, clustering was performed on *z*-scores where the *z*-score is the number of standard deviations away from the mean a signal. This can be expressed for a given pixel *x* as:

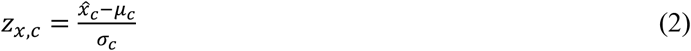

where *z*_*x,c*_ is the *z*-score for pixel *x* in channel *c*, and μ_*c*_ and *σ*_*c*_ are the overall mean and standard deviation of all pixels in channel *c*. This can be done to pixels with either non-scaled intensities (*x*_*c*_) or scaled intensities (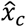) as in Eq1.

### LUMoS algorithm

We here present an unsupervised machine learning clustering method (Learning Unsupervised Means of Spectra, LUMoS) to learn the spectral signatures of each fluorophore and assign each pixel to the cluster whose spectral signature is closest. The process is referred to as “unsupervised” because no human intervention is required to label any pixels as belonging to a particular fluorophore, and the algorithm can identify features from raw images simply by looking at the pixels’ intensity values across all the detection channels. Specifically, a hard clustering method, *k*-means clustering, was used to separate mixed fluorophores unambiguously.

Pixels that are spatially close tend to belong to the same structure, and thus stained by the same fluorophore. To leverage this spatial information to improve the unmixing ability of LUMoS beyond the single-pixel level, a median filter (3×3×3 or 5×5×5) is first applied to the image before clustering. In order to preserve potentially meaningful variations in intensity in the raw image, the median filter is only applied at the clustering stage and the intensity output for each pixel is still taken from the raw image.

Given a set of observations **X**, containing *n* individual observations: *x*_1_, *x*_2_,…, *x*_*n*_, the objective of *k*-means is to partition all observations into *k* different clusters, **S** = {*S*_1_, *S*_2_,…, *S*_*k*_}, in a way that minimizes within-cluster variance. This can be expressed as

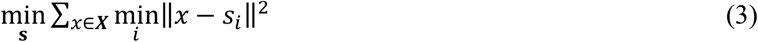

where *s*_*i*_ is the centroid of cluster *S*_*i*_. Unlike other applications where *k* is difficult to define and requires tuning to optimize, in our case, the number of clusters *k* is simply the total number of fluorophores plus 1 more cluster that represents the background. The cluster centroid resulting from this approach can be interpreted as the spectral signatures of each fluorophores. These are the spectral means alluded to in the name *Learning Unsupervised Means of Spectra (LUMoS)*.

The algorithm partitions the data into *k* clusters using Eq3 as a loss function. *K*-means approximates the solution to minimize the loss function by assigning data points to the class to whose centroid they are closest, and iteratively updating the centroid. Fig 1 details the steps taken in LUMoS. There are several algorithms for initializing the cluster centroids and we implemented the *k*-means++ initialization algorithm for its speed and convergence properties [36]. Briefly, *k*-means++ chooses the first cluster centroid at random from the input data points, and each subsequent cluster centroid is selected from the remaining data points with the probability inversely related to the distance from the closest appointed centroid. The algorithm converges when clusters do not change following one iteration. The maximum number of iterations allowed per replicate, *max_iter*, was set to 100 to limit run time. The iterative algorithm was applied *num_replicates* times and the replicate with the lowest cost was used in accordance with the loss function given in Eq3. All the unmixing performed in this paper used 10 replicates. The values of *num_replicates* and *max_iter* can be tuned, with more replicates and iterations yielding higher quality results but longer runtime.

**Fig 1.**
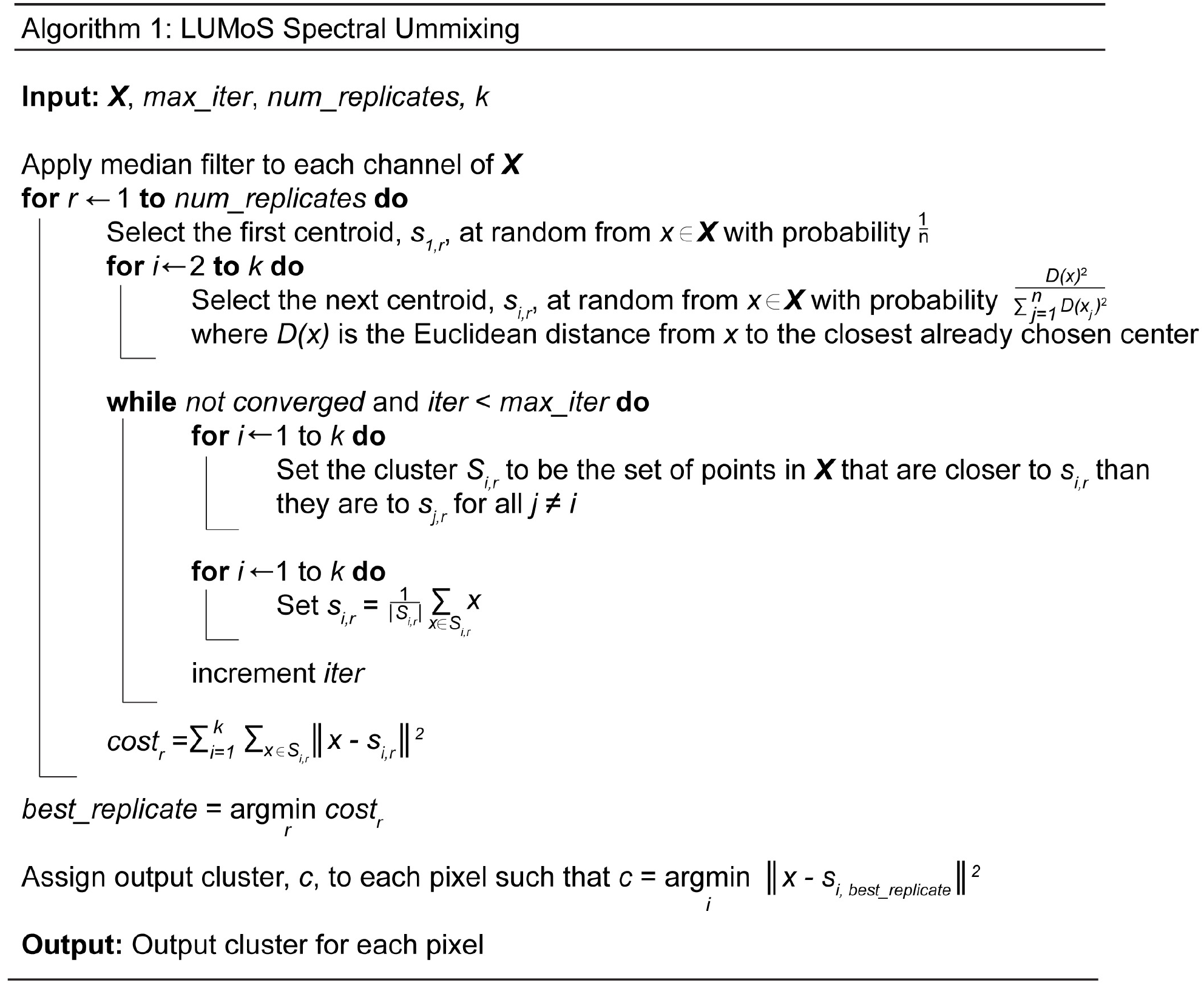
LUMoS spectral unmixing algorithm.

Once the algorithm converges, a new output image is created with *k* channels where each channel belongs to one cluster. In the output image, a pixel *x* assigned to one channel *c* is given the value of the highest intensity of that pixel among all the *C* input channels, and any pixel not belonging to channel *c* is assigned a value of 0:

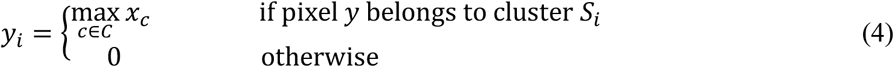

where *y*_*i*_ is the intensity of output pixel *y* in output channel *i*.

At its core, spectral unmixing is the task of decomposing mixed multichannel images into spectral signatures and abundances of each signature in each pixel [9,37,38]:

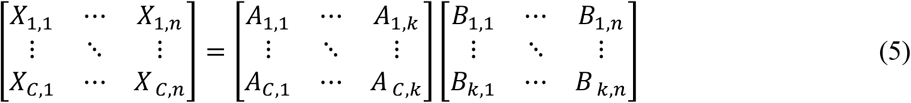

which may be simplified as: *X* = *AB*.

In Eq5, *X* is the observed fluorescence intensities of *n* pixels in *C* different spectral channels. The endmembers are the known fluorophores used to label the sample. *A* is a *C* × *k* matrix of the spectral signatures for each of the *k* fluorophores, in which each column is the recorded intensity of a fluorophore across the *C* detection channels. *B* is a *k* × *n* matrix containing the abundances of each fluorophore in each pixel. In LUMoS unmixing, *B* is obtained by scaling each pixel’s class label to the original intensity of that pixel as described in Eq4, which is based on a binary assumption that each pixel is occupied by only one fluorophore. Unlike other linear unmixing algorithms, LUMoS unmixes based on clustering rather than directly solving Eq5 with linear methods; because of this, LUMoS is different in that 1) the prior knowledge of fluorophore spectra (*A*) is not required to do the inversion of the equation and calculate the abundances (*B*), 2) it is not required that the number of fluorophores or endmembers (*k*) must be less than the number of detection channels (*C*), and 3) the abundances (*B*) are not the fractions of all endmembers, but are binary results assuming one endmember per pixel (Eq4).

### Synthetic data

In order to test the capabilities of LUMoS across a wide range of conditions, we generated synthetic data for unmixing. We assumed the hardware for the simulated imaging to be the same as our two-photon system with 2 two-photon lasers and 4 detection channels (blue: 420-460nm, green: 495-540nm, red: 575-630nm and far-red 645-685nm). For each simulated fluorophore, a theoretical emission spectrum was generated (Fig 2A). The intensity distribution was modeled as a Weibull distribution (Eq6) with *a* = 1.7 and *b* = 100 to reflect the tendency of a fluorochrome to have a long tail at the longer wavelength [37].

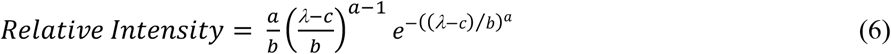

where *λ* is emission wavelength and c is a constant to shift the peak of the emission spectra for different synthetic fluorophores.

**Fig 2.**
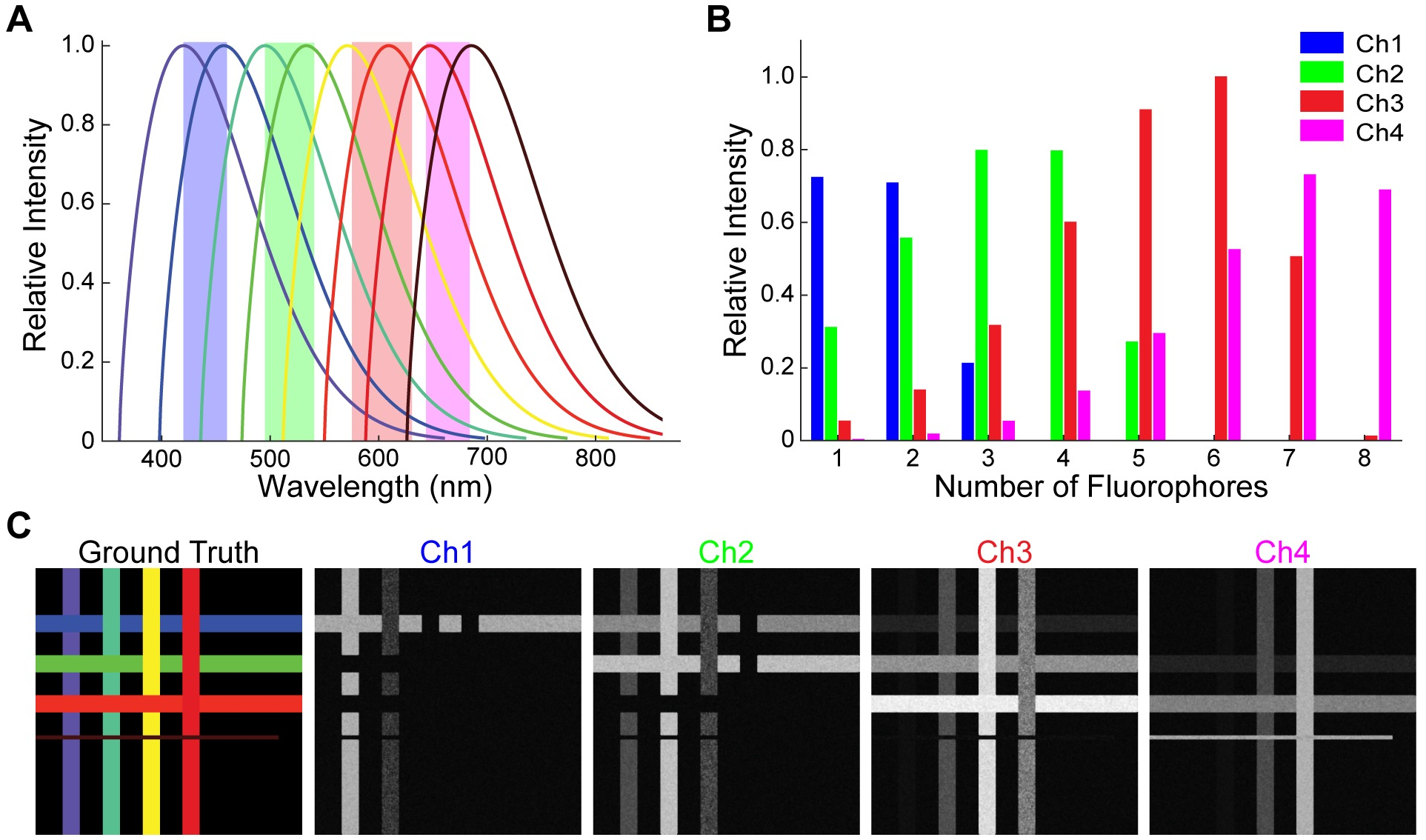
Synthetic data. (A) Synthetic emission spectra of 8 fluorophores. Bandwidth of the 4 detection channels were marked in shaded areas. For simplicity, the emission spectra of synthetic fluorophores were assumed to be the same Weibull distribution with the peaks of all fluorophores evenly distributed between 420nm and 685nm. (B) Spectral signatures of the 8 synthetic fluorophores in A. The intensity of each fluorophore was measured as the integrated area under the spectral curves in A. (C) Synthetic 2PLSM images based on the emission spectra in A. The ground truth image shows the 8 synthetic fluorophore expressing structures. Ch1-Ch4 images were the raw images from the 4 detection channels with an SNR of 10. Each fluorophore was synthetized to be expressed in a narrow band either vertically or horizontally. 7 of the 8 bands had the same area, while 1 small band (furthest red fluorophore) has an area 1/5^th^ (cluster size ratio 0.2) of the rest.

The emission peaks were evenly spaced between 420nm and 685nm so that all fluorophore peaks fell within the detection range of the microscope. We assumed all fluorophores were excited effectively, and their emission spectra peak at the same magnitude. Consistent spectral shapes and spacing represented an ideal case for easy simulations, but in reality, fluorophores usually have different shapes of spectra or even multiple peaks. To facilitate the generation of a synthetic image for an arbitrary number of fluorophores, a grid pattern was created where each successive fluorophore occupies an alternating vertical or horizontal strip (Fig 2C). For this synthetic data, the ideal situation where each pixel is occupied by only one fluorophore is assumed, to mimic the general biological staining assumptions without considering the nano-scale colocalizations caused by spatial resolution limitations. This pattern is overlaid onto a background with no fluorophores present. All synthetic images were 512×512 pixels. The length and width of the strip of the furthest red fluorophore was set variable while the rest of fluorophore expressing strips were fixed to be 512 pixels long and 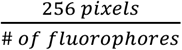 wide. This allowed us to measure the performance of LUMoS with unbalanced structure sizes. Cluster size ratio was the area of the minor fluorophore strip (furthest red) divided by the area of the major fluorophore strip.

Within each strip, all pixels belong to the same fluorophore but they all have slightly different emission spectra from the ideal value expressed in Eq6. Each pixel’s adjusted spectrum was shifted by a randomly selected wavelength with a standard deviation of 10nm to represent the variance present in real imaging. A four-channel representation of the pixel was then generated by integrating the emission spectrum within the bandpass of the detection channels (Fig 2B). For pixels with no fluorophore, a small background noise was added from a Gaussian distribution with a mean of 2 and standard deviation of 1. Additional Poisson noise was then applied to each channel to mimic the shot noise. At the end, the image was convolved with a Gaussian filter with a standard deviation of 0.5 and a 3×3 median filter to represent real-world diffusion effects.

Synthetic data unmixing performance was evaluated with the F1 score between the LUMoS output and the ground truth image (Fig 2C):

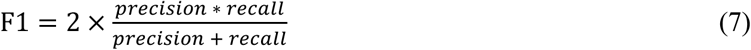

where 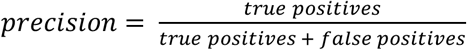, and 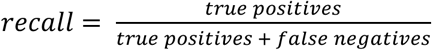.

## Results

### N_fluorophores_ = N_detectors_

First, we started with a simple case in which there was same number of fluorophores as imaging channels. BPAE cells with nuclei stained with DAPI, F-actin labeled with AlexaFluor488 (AF488), and mitochondria labeled with MitoTracker Red were imaged using 780nm laser [39,40] to excite all three fluorophores (Fig 3A). Due to the long tail of the DAPI emission spectrum (Fig 3C), F-actin signals in the green channel were contaminated by the nuclei signals (Fig 3A). DAPI had strong signals in both blue and green channels, while AF488 and MitoTracker Red were distinct in green and red channels respectively. Therefore, each fluorophore had a unique distribution of intensity across channels—"spectral signature”, calculated as the intensity of the pixels in one LUMoS cluster detected by each channel in the raw image (Fig 3D). LUMoS was able to group pixels with similar spectral signatures into the same cluster and re-assign each pixel into the correct fluorophore cluster. As only the blue and green channels had bleed-through issues, we applied LUMoS unmixing only on these two channel images, and produced 3 output channels (DAPI, AF488, and background). After the unmixing procedure, the spectral overlap of the DAPI and AF488 was corrected, and the unmixed images now represent the abundance of each of the fluorophores (Fig 3B, the 3D unmixing results were shown in S2 Movie).

**Fig 3.**
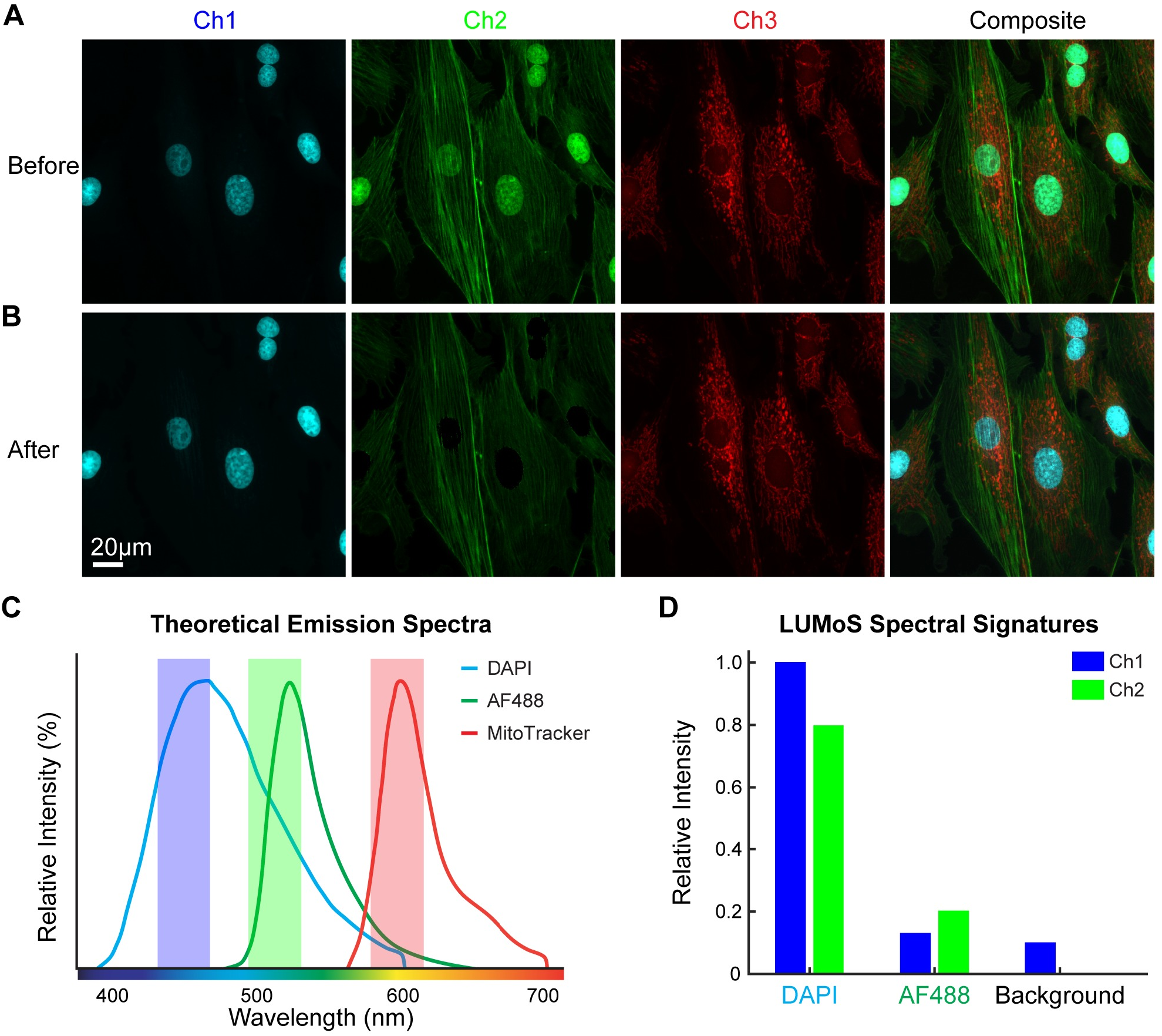
LUMoS unmixing of BPAE cells with channels bleed-through. (A) BPAE cell stained with DAPI in nuclei, AF488 in actin, and MitoTracker Red in mitochondria, and imaged with 2PLSM. Images shown were 2D maximum intensity projections of 3D z-stacks. The green channel (Ch2) had a mix of actin and nuclei with the DAPI signals bleeding into the AF488. (B) LUMoS unmixing results of the mixed images in A. Only Ch1 and Ch2 images were used for separation. Note the clear separation of the nuclei from the green channel after unmixing. Background pixels were removed. (C) The theoretical emission spectra of DAPI, AF488 and MitoTracker Red. The filter bandwidths were plotted as shaded areas. Note the long tail of the DAPI spectrum blending into the green channel. (D) The relative intensities of the LUMoS unmixed DAPI and AF488 pixel clusters detected by the green and blue channels. Background was separated as an additional cluster with relatively low intensity in both channels.

### N_fluorophores_ > N_detectors_

The two-photon excitation spectrum of a fluorophore is usually broader than the one-photon spectra and may have multiple peaks [7,41], making it possible to just use one or two two-photon laser lines to excite multiple fluorophores simultaneously, which is both time and cost efficient. On the other hand, simultaneous excitation also leads to the issue of channel cross-talk which limits the number of detection channels to usually less than 4 for two-photon microscopy. This makes the ability to image more fluorophores than detectors crucial for many applications. As the LUMoS method has no intrinsic requirement that the number of channels be at least equal to number of fluorophores, we next ascertained the limit of our method by imaging more colors simultaneously without modifying the imaging hardware.

To test the performance of LUMoS on a sample with more fluorophores than detectors, we first imaged mixed beads with 5 different fluorophores: LY (Light Yellow dye from Spherotech Inc.), FITC, PE, Purple (Purple dye from Spherotech Inc.), and APC (Fig 4A). The theoretical emission spectra are shown in Fig 4C. Simultaneous two-photon excitations at 800nm (MaiTai laser) and 1100nm (InsightX3 laser) were used to excite all fluorophores [39]. Because of the significant emission spectra overlaps of LY and FITC in the green channel, PE and Purple in the red channel, and PE, Purple and APC in the far-red channel, the raw images collected by the 4 detectors (Fig 4A) showed many beads appearing in more than one channel (examples are indicated by white arrows in Fig 4A). The spectral signatures of those fluorophores (Fig 4D) were consistent with the emission spectra information in each channel, which demonstrated the uniqueness of each fluorophore’s intensity distribution across the 4 detectors. We therefore applied LUMoS with 6 clusters to the raw 5-color beads images. The algorithm generated 6 new images in which 1 image included all background pixels and the other 5 images each represented one single fluorophore. We removed the background to get the clean unmixed outputs (Fig 4B). The algorithm performed well to fully separate out the 5-color beads with individual beads belonging only to a single output channel.

**Fig 4.**
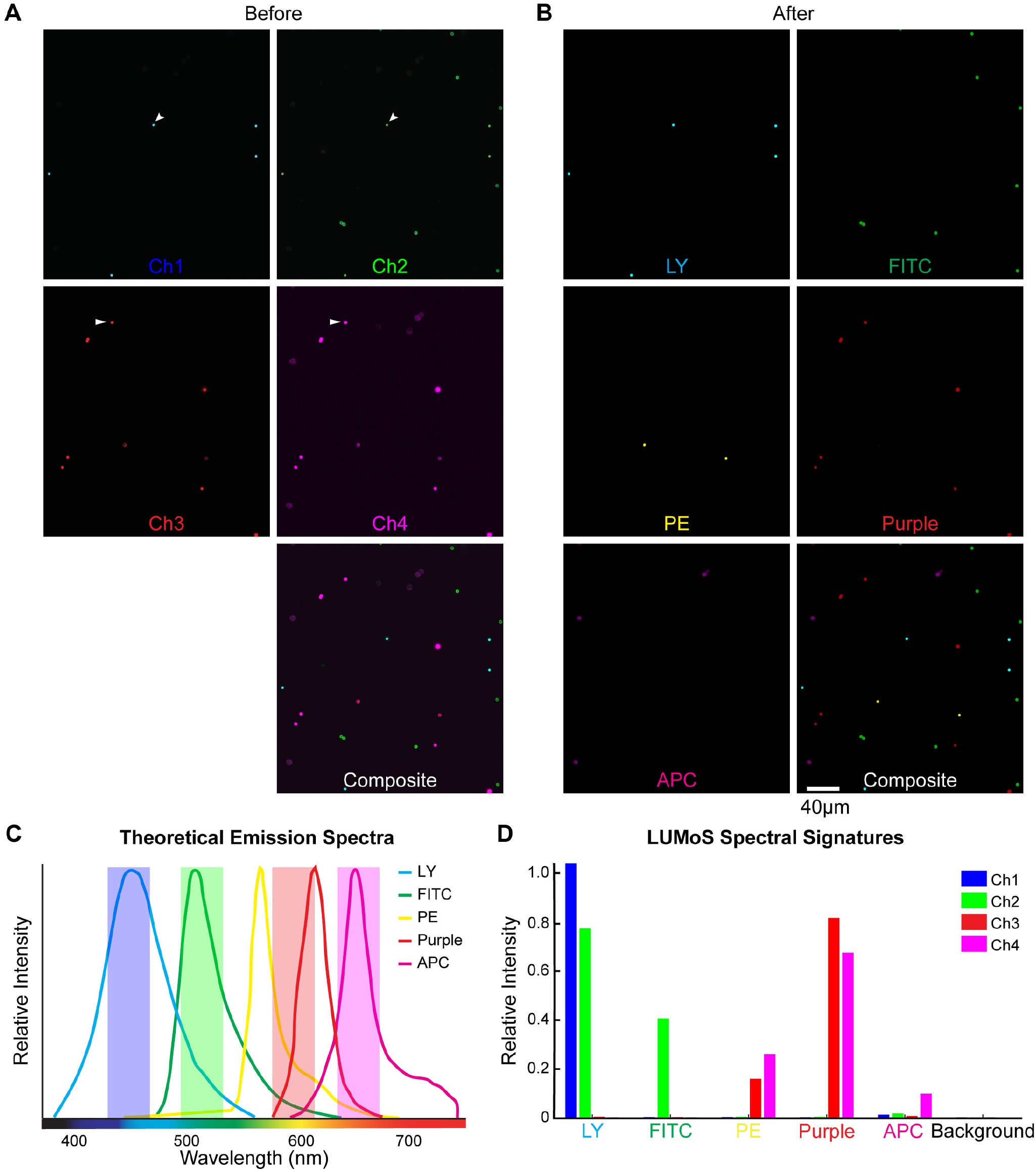
LUMoS unmixing of 5-color beads mixed in 4 detection channels. (A) Mixed beads stained with Light Yellow (LY), FITC, PE, Purple, and APC imaged with 2PLSM. LY and Purple are special dyes by Spherotech. It was unable to resolve 5 colors with 4 detectors, and there were also beads shown in more than one channels. Two examples are pointed out by white arrows. (B) The images shown in A after processing through LUMoS. The first five images show the fluorescence of the five different beads separated from the 4 detection channels by the LUMoS and the last image is the composite showing all five beads as clearly separated objects. (C) Theoretical emission spectra of the 5 fluorophores. LY and Purple spectra were obtained from Spherotech, and FITC, PE and APC were obtained from online spectra-viewer. There were significant overlaps of all the 5 fluorophores. (D) The relative intensity of the pixels of each separated fluorophore in the 4 channels. Each fluorophore was represented with a unique spectral signature. Background pixels formed one additional cluster with low pixel intensities in all the channels.

Commonly used dyes differ not only in their emission spectra but also their excitation spectra. The differences in excitation efficiency offers additional features for LUMoS to better separate out more fluorophores. In the next example, we used sequential scan by alternating two-photon excitations at 840nm (MaiTai laser) and 1050nm (InsightX3 laser) to visualize 6 compartments with distinct labels in one single cell (Colorful Cell). Human embryonic kidney cells (HEK293) were transiently transfected with a plasmid that encodes differentially localized fluorescent proteins. The cells express tagBFP in nucleus, Cerulean in cell membrane, AzamiGreen in mitochondria, Citrine in Golgi body, mCherry in endoplasmic reticulum (ER), and iRFP670 in peroxisome (Fig 5A). Cerulean, AzamiGreen and Citrine all have significant emissions in the green channel (Fig 5B), but they are excited at different efficiencies under 840nm and 1050nm [7,39], making it possible to distinguish them with the spectral signatures by collecting the green channel twice with the two excitations (Fig 5E). The 2PLSM excitation/emission setup is shown in Fig 5B. All the organelles were ambiguously mixed in the raw images especially in the green, red and far-red channels (Fig 5C). We assigned 7 clusters to the LUMoS algorithm to separate out the 6 fluorophores and background from the original 5-channel images. The algorithm reliably separated the raw data into 6 components that corresponded to the 6 organelles (Fig 5D) based on their shapes and locations inside the cell by comparing to the cell structure schematic (Fig 5A). The 3D unmixing results were shown in S3 Movie.

**Fig 5.**
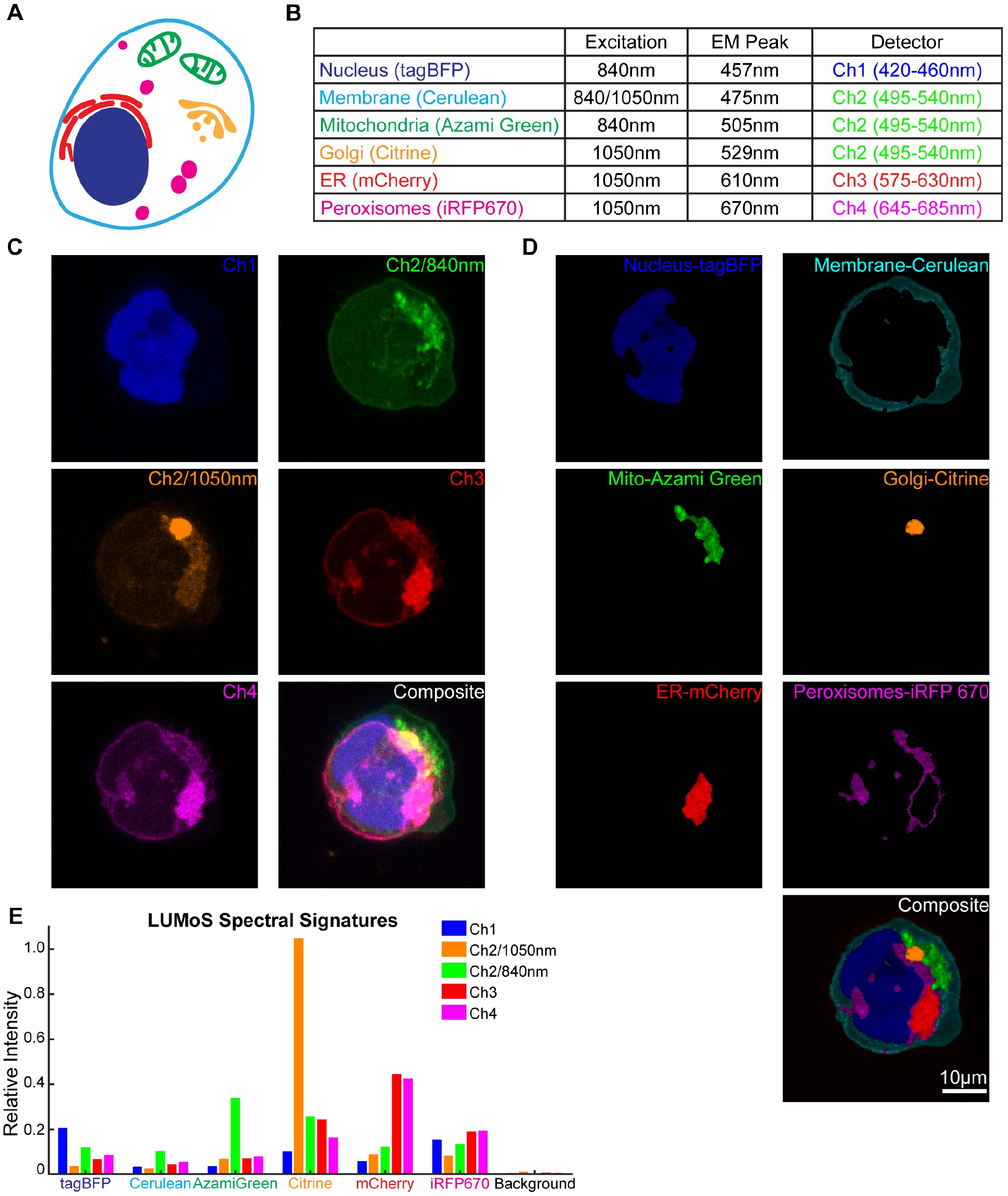
LUMoS unmixing of the Colorful Cell expressing 6 colors. (A) Schematic of the Colorful Cell expressing BFP in nucleus, Cerulean in cell membrane, AzamiGreen in mitochondria, Citrine in Golgi bodies, mCherry in endoplasmic reticulum, and iRFP 670 in peroxisomes. (B) The 2PLSM system excitation and emission setups for imaging the Colorful Cell. 840nm and 1050nm sequential scan was conducted for the green channel (Ch2). Cerulean, AzamiGreen, and Citrine all emit significantly in the green channel. Cerulean can be excited well at both 840nm and 1050nm. AzamiGreen had more excitation at 1050nm, while Citrine excited better at 840nm. (C) The raw 2PLSM images of the Colorful Cell in the 4 channels with 2 excitation wavelengths for the green channel (Ch2). All the fluorophores were mixed in the detection channels which made it difficult to reveal individual organelles. Images were maximum intensity projections of 3D z-stacks. (D) LUMoS separation results of the images in C. 6 distinct organelles were separated into individual images and a composite image of all 6 colors is shown on the bottom. Signals from background pixels were removed. (E) The relative intensities of each separated fluorophore by LUMoS in the detection channels.

### Colocalization analysis

Unlike linear unmixing [10,18], one of the major assumptions of the LUMoS algorithm is that one pixel is uniquely labeled with one fluorophore, which is advantageous in the way that it provides unambiguous results especially in biological imaging (examples in Figs 3–5). However, in biology, one structure is often labeled with more than one fluorophores for colocalization studies. The structures with colocalized labeling will exhibit a distinct spectral signature, which is usually the combination of, but is different from, the individual fluorophore’s spectrum. By leveraging this, LUMoS is able to treat the colocalized fluorophores as an additional cluster, and separate out the pixels with colocalization.

To demonstrate the flexibility of LUMoS to unmix and analyze images with colocalized labels, CD28 virus labeled with Cerulean or YFP was used to transduce T cells either separately or together. The T cells were then mixed with non-labeled antigen-presenting cells (APCs) to form conjugations [34]. Cerulean or YFP was recruited and concentrated at the T-cell and APC contact sites. When T cells were transduced by Cerulean or YFP virus separately, the Cerulean and YFP were detected by the CFP and YFP channels respectively without bleed-through (S4 Fig). When T cells were transduced by the mix of Cerulean and YFP viruses, some T cells expressed both Cerulean and YFP, while some only expressed one of them (Fig 6A). LUMoS was able to separate the raw images into Cerulean-only, YFP-only, and Cerulean+YFP colocalized channels (Fig 6B), by identifying distinct spectral signatures (Fig 6C). The calculated Mander’s colocalization coefficients were 44.2% (M_Cerulean_) and 38.2% (M_YFP_) [42]. In addition, although APCs were not labeled, they showed some autofluorescence in the raw images (Fig 6A indicated by white arrows, and S4 Fig). Similar as background noise (S6 FigD), autofluorescence was also identified and separated out by LUMoS (Fig 6B). The 3D unmixing results were shown in S5 Movie.

**Fig 6.**
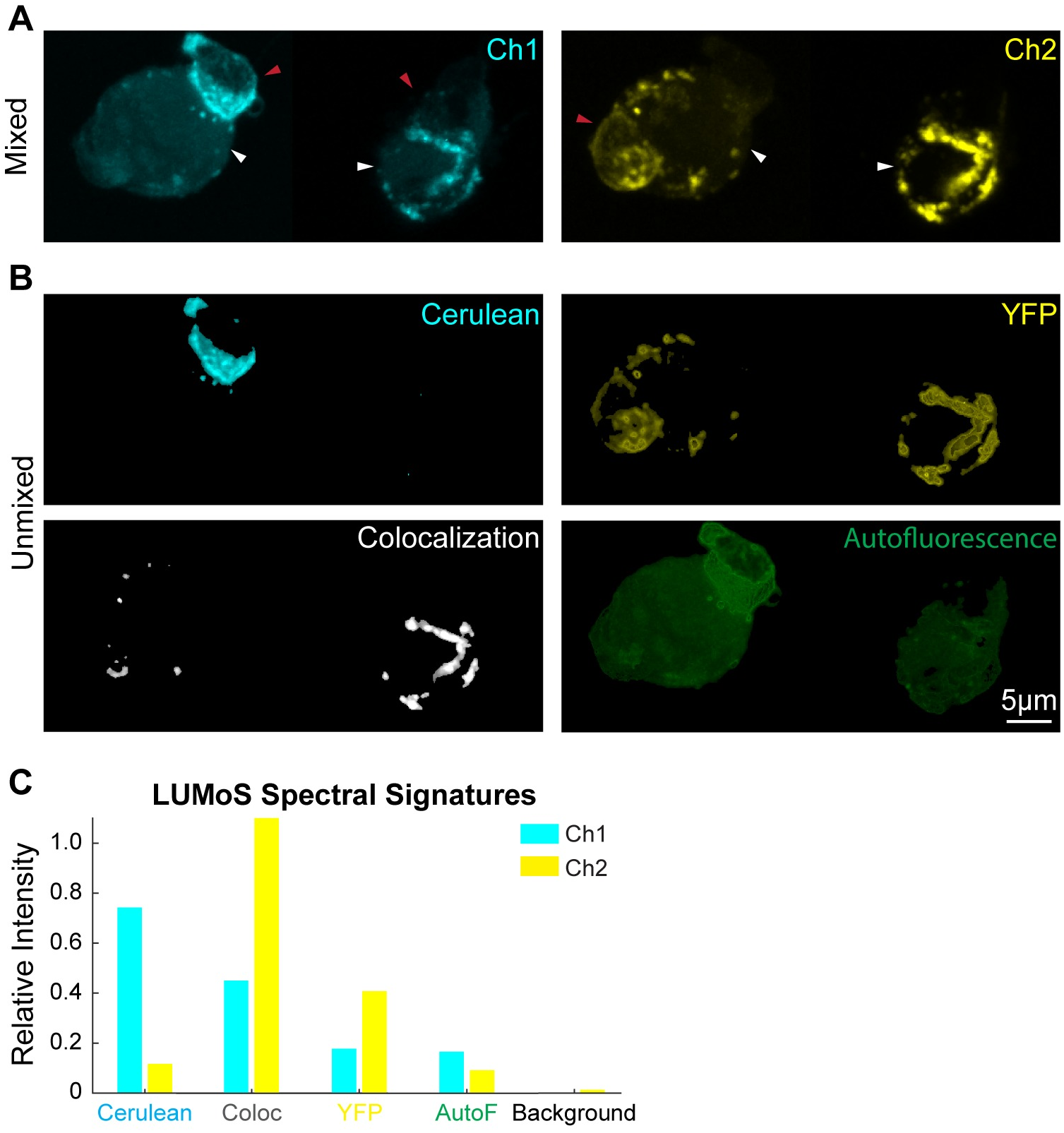
LUMoS unmixing for colocalization analysis and autofluorescence removal. (A) The raw 2-channel 2PLSM images of T cells expressing Cerulean, YFP, or colocalized Cerulean and YFP. APCs are larger cells (pointed out by white arrows) than T cells (pointed out by red arrows), and APCs are non-labeled but autofluorescent. The fluorophores were concentrated at the conjugation sites between T cells and APCs. The images were z-projections of slices 6 to 17 of 3D z-stack images (S5 Movie). The left and right cells were imaged by two acquisitions and stitched, but with the same imaging conditions. (B) LUMoS separation results of the images in A. Autofluorescence and colocalization were split into separate channels while keeping pure Cerulean and YFP signals in their own channels. Signals from background pixels were separated and removed (S6 FigD). (C) The spectral signatures of each structures produced by LUMoS. Background and autofluorescence (AutoF) were identified as additional pixel groups with distinct signatures. Colocalization (Coloc) spots were separated out due to its different spectral signature from the Cerulean-only and YFP-only groups.

### Background and autofluorescence removal

Most spectral unmixing tools [19,25,26,43] cannot distinguish background noise from real signals, while background removal is usually an essential prerequisite before unmixing to remove any signal not originating from the targeting signals [37]. Usually, if significant background noise exists, a simple math subtraction with a specific pixel threshold measured from non-structure background is performed, which can have the undesirable effect of removing real signals. The LUMoS method does not rely on a fixed numerical background subtraction, but rather the background is treated as a separate cluster with a spectral signature different from fluorophore expressing signals, so that background noise can be separated (S6 Fig) and removed from the sample signals (Figs 3D, 4D, 5E, 6B). Therefore, the outputs of LUMoS are cleaned in the way that they are both spectral unmixed and background removed.

Autofluorescence is another a common but usually undesired signal in fluorescence microscopy in which regions with no label are fluorescent, often with higher intensity and broader emission spectrum than individual fluorophores [43]. Autofluorescence can come from some extracellular components or some cell types [44]. Non-negative matrix factorization (NMF) is one spectral unmixing method that has been successfully applied for autofluorescence removal [22–24]. We here also demonstrated the unmixing performance of LUMoS when autofluorescence exists. In Fig 6A, the APCs in the sample were not stained but were autofluorescent. Similar to background, autofluorescence can be treated as an additional cluster if it exhibits a distinct spectral signature among all the fluorophores in the sample (Fig 6C). LUMoS was able to detect and remove autofluorescence in the image (Fig 6B). However, if the emission spectrum of autofluorescence is similar to other fluorophores in the image, the autofluorescence may be hard to separate out, so additional detection channels may be helpful to unmix the images in such cases.

### Synthetic data

Lastly, we sought to test the limitations of LUMoS spectral unmixing by understanding the smallest structure size which can be detected, the maximum number of fluorophores the algorithm can separate, and the minimal quality of the input image that is required. As it is impractical to prepare a real-world biological sample with arbitrarily many fluorophores and precisely control both the size of a stained structure and the image SNR, we used synthetic images with those conditions computationally manipulated (Fig 2A-C). The synthetic data also provides us a ground truth to evaluate the performance of the algorithm.

#### Cluster size

As LUMoS is a *k*-means clustering based method, the algorithm assumes similar amount of data points in each cluster, and can disregard small but real clusters in order to minimize the total loss function [32]. This may be problematic when one fluorophore expressing structure is represented by significantly fewer pixels than the other structures, in which case the algorithm will misclassify the pixels belonging to a more abundant fluorophore to the minor structure, leading to an unmixing failure. Therefore, we first tested the robustness of the algorithm by changing the size of one fluorophore expressing structure while keeping the size of the rest of structures fixed. The number of fluorophores and SNR were fixed at 8 and 10 respectively. F1 score was used as the evaluation metric as it can detect when the algorithm starts to erroneously combine fluorophores. The F1 score of the smallest cluster was used because the smallest cluster is inherently the most difficult for LUMoS to recognize and represents the worst-case scenario. Performance was monitored by setting the threshold for successfully unmixed samples at an F1 score of 0.9 or higher on the smallest cluster. The F1 score for the smallest cluster dropped off sharply when decreasing the cluster size ratio to below 0.01 (Fig 7, left), because at the tipping point, one part of a larger cluster was merged with the smallest cluster as the algorithm prioritized the improvements to other dominant clusters. This happened to all of the pixels in a small cluster at once so the drop off in accuracy was sudden. This can make LUMoS vulnerable when one fluorophore is expressed in much smaller structures than the rest.

**Fig 7.**
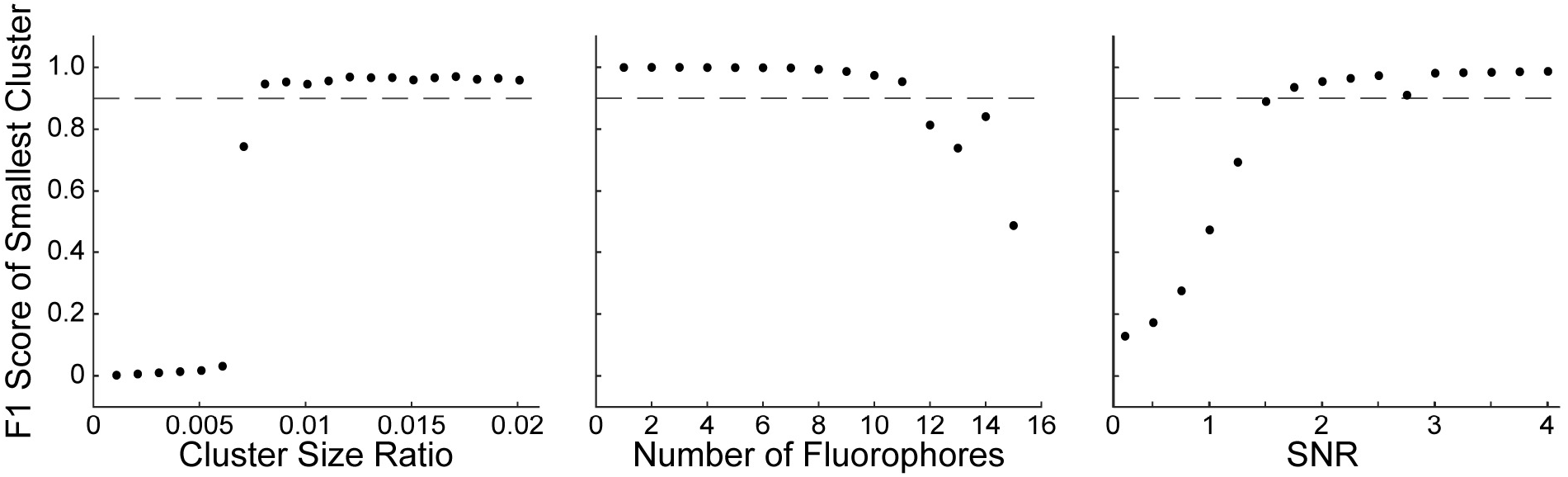
Simulation tests of the performance of LUMoS. *Left*, the performance of LUMoS with unbalanced structure size. The number of fluorophores was fixed at 8 and SNR at 10. *Middle*, the performance of LUMoS with increasing number of fluorophores. The cluster size ratio was fixed at 0.2 and SNR at 10. *Right*, the performance of LUMoS with SNR varying. The cluster size ratio was fixed at 0.2 and number of fluorophores at 8. Results of 10 simulations were averaged to obtain all the final results.

#### Number of fluorophores

The natural questions that follow from the analysis are: what is the maximum number of fluorophores that can be separated, and what is the extent of spectral overlap that can be successfully unmixed. To address these questions, we challenged the algorithm by increasing the number of fluorophores until it failed (Fig 7, middle). The cluster size and SNR were held constant at 0.2 and 10 respectively, and F1 score of the smallest cluster was measured. All fluorophores were assumed: 1) to be effectively excited, 2) to have the same shape and intensity scale of emission spectra with a tail into the longer wavelength, 3) to have emission peaks evenly distributed between 420nm and 685nm (Fig 2A). To mimic the variations in real-world imaging, the spectra of pixels belonging to one fluorophore were randomly shifted with a standard deviation of 10nm (S7 FigA). The imaging hardware was assumed to be the same as our system. LUMoS’s performance was very stable until the number of fluorophores reached 12 (Fig 7, middle). At this point, the mean emission peaks were 37nm apart and there was 72% emission spectra overlap. We also tested the performance of LUMoS on synthetic images of two fluorophores with varying differences in emission peaks (S7 FigB). The peak of the lower wavelength fluorophore was fixed while the peak of the higher wavelength fluorophore was varied to evaluate performance at different peak distances. Depending on where in the range of detectors they fell, the peaks of two fluorophores could be 10-15nm apart and the fluorophores could still be separated by LUMoS. This 10-15nm peak distance represents an 88-92% overlap in ideal emission spectra, which is very close to the standard deviation of 10nm with which each pixel’s individual emission peak was sampled (S7 FigA). This variance in emission spectra from pixel to pixel is a key limiting factor in how similar the emission spectra of two fluorophores can be while maintaining separability with LUMoS. In real-world cases, the fluorochromes in a biology sample will not be as ideal as the simulated scenario. Careful selections of dyes with relatively separated emission spectra are always desired to gain the best unmixing results.

#### Signal-to-noise ratio

All spectral unmixing methods require a good image quality. LUMoS is a pixel-based method which makes it susceptible to any noise detected at the same time with real signals. Therefore, we tested the performance of LUMoS for unmixing images with different SNRs (Fig 7, right). The cluster size ratio and number of fluorophores were fixed at 0.2 and 8, while F1 score of the smallest cluster was evaluated at different SNRs. The simulated data showed that LUMoS was very robust when the SNR was above 2. For images with SNRs around that level or lower, LUMoS will likely have low performance on the raw data. Even with ideal spectral signatures, any pixel-level unmixing techniques such as LUMoS will fail when the observed spectral signature is contaminated by high noise. In cases where the image to be unmixed is prohibitively noisy, denoising pre-processing techniques or an unmixing method that can take spatial information into account may be desired.

### ImageJ PlugIn

Although many spectral unmixing algorithms have been published, so far, easy-to-use open source tool options are still limited to biologists. Walter published an ImageJ/Fiji spectral unmixing plugin [45] based on linear unmixing, which requires either a reference image with well-separated structures or a separate preparation of reference samples for each fluorophore. Those requirements are usually hard to achieve, and the PlugIn also involves laborious and time-consuming manual ROI labeling. Another unmixing PlugIn available is based on spectral deconvolution [46], but also requires ROI selections of areas with only one type of fluorophore. We here developed an ImageJ/Fiji PlugIn [47,48] of the LUMoS algorithm to facilitate the easy implementation of this flexible method for spectral unmixing, background removal, and colocalization analysis. No ROI selections, spectra information, or single stain of samples are required. The only input parameter is the number of fluorophores in the sample. The PlugIn is available from the authors or through ImageJ PlugIn Repository. Detailed user guides are provided on our website (https://www.urmc.rochester.edu/research/multiphoton/image-analysis/spectral-unmixing.aspx).

## Discussion

Over the past decade, a wide variety of high-performance fluorophores have been developed [49,50]. These reagents exhibit a broad range of physical and spectral properties [51], are capable of targeting proteins or peptides in living or fixed cells [40], and can also be used as indicators of biological dynamics [52]. Combining two or more fluorescent probes offers significantly a higher level of information [25,53,54], but may also lead to signal crossover [9]. Current spectral unmixing tools solve this problem to some extent, but their applicability is usually limited. In this paper, we suggested and experimentally examined an approach by using *k*-means clustering based unsupervised machine learning as a more flexible alternative to separating mixed images blindly.

There are two major issues with current unmixing tools available to biologists which have highly restricted the spectral resolutions that can be achieved by fluorescence microscopy especially the 2PLSM. Firstly, unmixing methods based on linear inversion calculations, such as linear unmixing [9,11,12,37,55], spectral deconvolution [25,46] and similarity unmixing [26], rely heavily on the cumbersome pre-measurements of emission spectra either through separately recording the spectra of all fluorochromes [26] or manually selecting ROIs with pure labels in the image [9]. Background and autofluorescence, if present, also need to be defined spectrally and treated as additional spectra [11,55], which are even harder to measure or estimate. LUMoS, as it does not directly calculate the abundances of fluorophores, is a completely “blind” unmixing process, and is therefore, much easier to implement and free from those restrictions of acquisition conditions. When background and autofluorescence are present in the sample, additional clusters could be added, and those undesired signals could be separated and removed (Fig 6). Secondly, linear unmixing, Non-negative Matrix Factorization (NMF) [20,56], deconvolution, and Principle Component Analysis (PCA) [57] all require determined (N_fluorophores_=N_channels_) or over-determined (N_fluorophores_<N_channels_) image acquisition systems, greatly restricting the total number of fluorophores that can be imaged by the hardware design. Although Independent Component Analysis (ICA) does not intrinsically require less fluorophores than detectors, its success for spectral unmixing in fluorescence microscopy has been limited to relatively few independent sources which are usually same or fewer than the number of detectors [58–60]. As LUMoS can be set to create an arbitrary number of clusters for an image, it can be used in under-determined situations (N_fluorophores_>N_channels_) for expanding the capabilities of an imaging system (Figs 4 and 5). Moreover, as the readout noise increases with the number of detection channels used [37,58], LUMoS can achieve the high quality unmixing results with as few channels as possible to minimize the readout noise.

Similar but more complicated clustering based methods have been introduced and developed in the field of satellite imaging [29,61,62]. Remote sensing image unmixing is similar to fluorescence image unmixing in many ways, and many unmixing ideas commonly used for microscopy imaging were initially introduced in remote sensing [37]. The ultimate goal of both imaging modalities’ unmixing is to decompose the spectral signature of mixed signals into a set of endmembers and corresponding abundances [38,63]. However, the uniqueness of fluorescence microscopy makes its spectral unmixing task different from remote sensing. First and foremost, the number and type of fluorophores (endmembers) are known in advance in microscopy, which offers a great advantage and simplicity of using clustering algorithms such as *k*-means for fluorescence image unmixing. Most of the time, the first step of remote sensing image unmixing is to determine endmember [38,64], and many of the advanced unmixing algorithms have been focused on how to better estimate the number and characteristics of endmembers, such as adaptive possibilistic clustering [62] and neural network autoencoder [65]. Second, due to the chemical mixtures of landscape objects, the abundance of one pixel from a satellite image normally comprises fractions of each endmembers, thus remote sensing image unmixing methods output abundances for each pixel as fractions of different chemical components [38,63]. However, in fluorescence microscopy, biologists usually assume a distinct labeling of a structure by one specific fluorophore, unless colocalized labeling was designed. The goal of fluorescence image unmixing is more towards unambiguously distinguishing each labeled structure rather than decomposing each pixel into many different chemical components. Therefore, using classification based hard clustering, such as LUMoS, by assuming one pixel per fluorophore is more appropriate in the field of fluorescence imaging and the results of which are more interpretable for biologists. Third, remote sensing images have hundreds of spectral bands which is usually much more than the number of endmembers, making linear algebra based unmixing methods, such as linear unmixing, NMF, and deconvolution, better suited [38,63,64]. Because fluorescence microscopes have much fewer detectors (usually ≤4), many unmixing methods applied for remote sensing are insufficient for fluorescence imaging with potentially more fluorophores than detectors. In considerations of those features of fluorescence imaging, we applied *k*-means clustering as a simple, easy-to-use, and flexible method for microscopy image unmixing.

The implications of *k*-means clustering are usually limited by the difficulties in choosing an optimal number of clusters, “*k*” [32,66]. However, in the case of fluorescence microscopy, “*k*” is known and determined by the number of fluorochromes used, making *k*-means clustering a well-suited method for spectral unmixing. Usually, the “*k*” is set to be the total number of fluorophores plus one (considering the background noise) (examples in Figs 3–5). When special circumstances happen, options are available to optimize the “*k*” to tailor LUMoS for different cases. For example, when there are known colocalization labeling or autofluorescence structures (Fig 6), additional clusters could be added by considering colocalization and autofluorescence as distinct “fluorophores”. When applying LUMoS, carefully examining the image data to better determine “*k*” in advance may improve the unmixing results.

There are also limitations of our algorithm, especially when unique circumstances are associated with the imaging data. As demonstrated in the simulations, our approach may cease to be useful when it misclassifies a significant portion of the pixels belonging to a fluorophore of interest. This can occur when there are relatively unbalanced structure sizes, significantly overlapping emission spectra, and a low SNR. Additionally, although considering the information of nearby pixels by using a median filter, LUMoS still does not take any spatial information at biological structure level into account so its clustering ability is limited to classifying individual pixels rather than whole structures as some other methods attempt [67,68]. LUMoS specifically assumes the abundance of each fluorophores is binary at pixel level, which produces unambiguous classification of individual fluorophores. If there is colocalization at structure scale, for example one structure labeled with more than one fluorophore, the colocalization group can be treated as an additional cluster to be separated and analyzed (Fig 6). However, implicit in our unmixing algorithm is the assumption that a pixel represents an exclusive single label without considering nano-scale colocalization due to the imaging spatial resolution limitations. This assumption is valid for spatially well-dispersed fluorescent structures relative to the imaging resolution, but may not hold when two labeled structures are contacting or too close to each other. We expect future improvements by adding the options of fuzzy clustering [69,70] or overlapping *k*-means [71] to extend the flexibility of LUMoS when there are nano-scale colocalization considerations.

In conclusion, we presented a blind and flexible tool for fluorescence image spectral unmixing — LUMoS. Both experimental and synthetic results demonstrated its ability to robustly separate mixed fluorophores in terms of the quality of results and ability to converge in a variety of scenarios. The LUMoS method has also greatly expanded the fluorophore options beyond the number limit of detectors and excitation lasers. These qualities make LUMoS a simple, general, and reliable spectral unmixing approach to quickly apply to any fluorescence images. Last but not least, an optimal strategy for spectral unmixing should always combine image processing algorithms with careful dye selections and rigorous image acquisitions. LUMoS can be coupled with spectral imaging or other hardware designs to yield excellent multi-color imaging results, and will offer new avenues for understanding the complex biological organizations.

## Acknowledgement

We thank the Multiphoton Research Core Facility at University of Rochester Medical Center for providing the two-photon microscope system. We also thank Matthew Cochran at the Flow Cytometry Shared Resource at University of Rochester Medical Center for generously providing some of the fluorescence particles used in the study.

## Supporting information

**S1 Fig.**
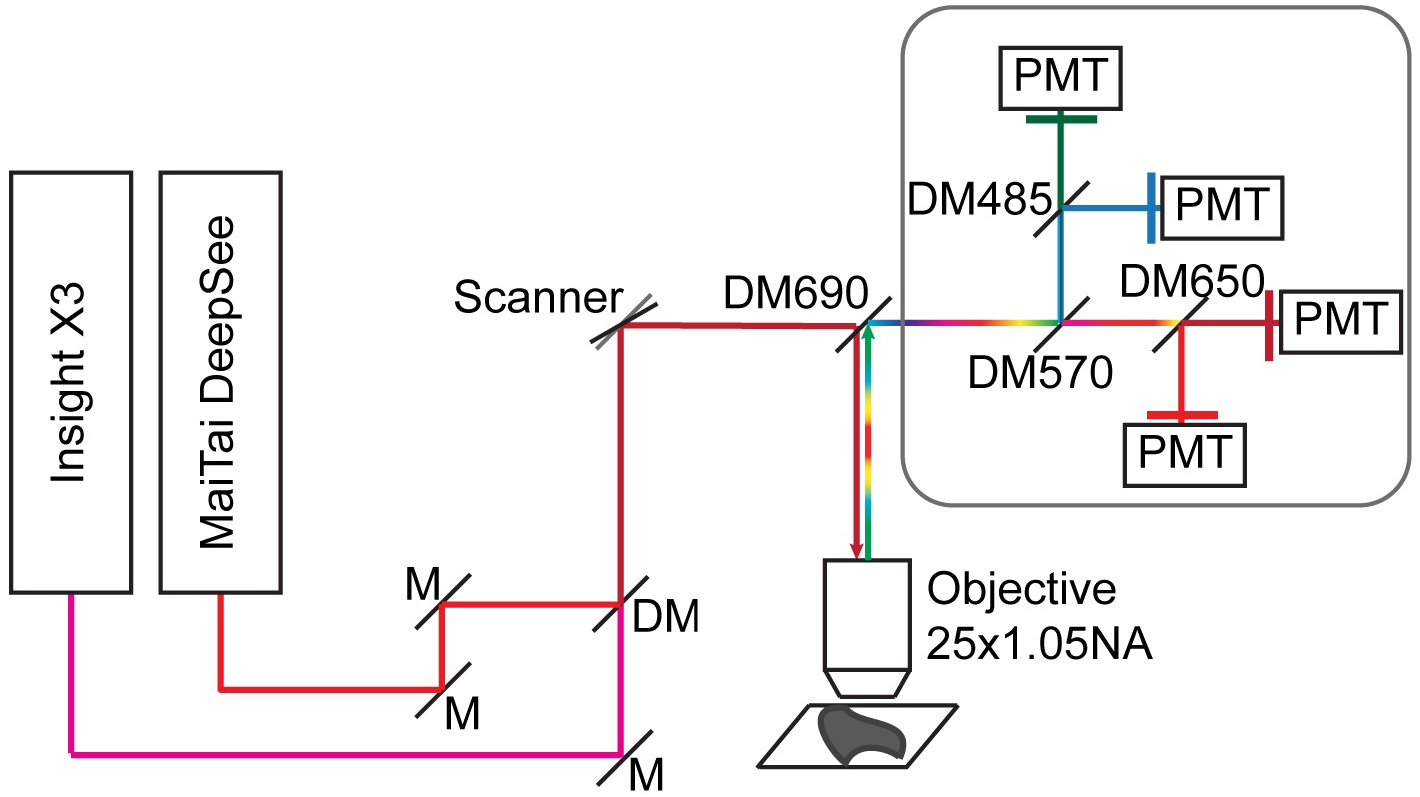
Two-photon system schematic. The system (Olympus FVMPE-RS) was equipped with two two-photon lasers and four PMTs. 25× water immersion objective was used. M: mirror, DM: dichroic mirror, Scanner: galvanometer scanner, PMT: photomultiplier tube. The Blue/Green (420-460nm/495-540nm), and Red/fRed (575-630nm/645-685nm) filter cubes setup is shown.

**S2 Movie. BPAE cells 3D image unmixing results.**

Z-stack of BPAE cells 2PLSM images shown in Fig 3. *Left*, raw image. *Right*, LUMoS unmixed image.

**S3 Movie. Colorful Cell 3D image unmixing results.**

Z-stack of Colorful Cell cells 2PLSM images shown in Fig 5. *Left*, raw image. *Right*, LUMoS unmixed image.

**S4 Fig.**
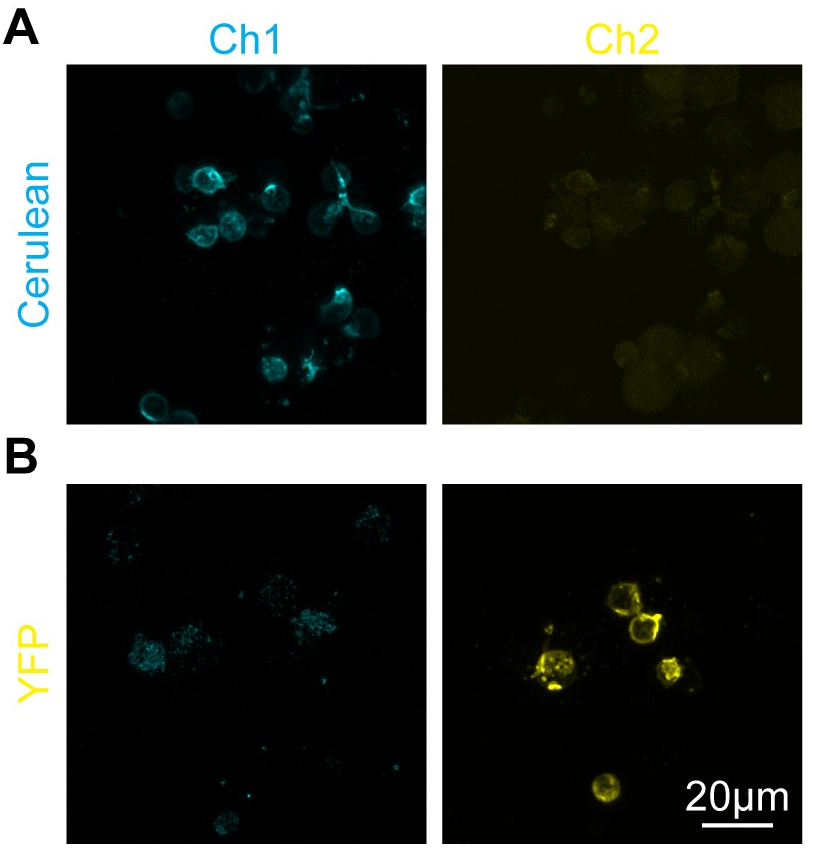
Single stained cell preparations of the colocalization example. T cells only transduced with Cerulean or YFP virus mixed with APCs and imaged with the same conditions as in Fig 6. (A) Cells transduced with Cerulean expressing virus. Cerulean signals showed only in CFP channel (Ch1). (B) Cells transduced with YFP virus. YFP signals were only detected by the YFP channel (Ch2). There was no cross-talk between CFP and YFP channels. APCs showed weak autofluorescence.

**S5 Movie. Cerulean and YFP colocalization 3D image unmixing results.**

Z-stack of T cells transduced with Cerulean and YFP virus shown in Fig 6. *Top*, raw image. *Bottom*, LUMoS unmixed image.

**S6 Fig.**
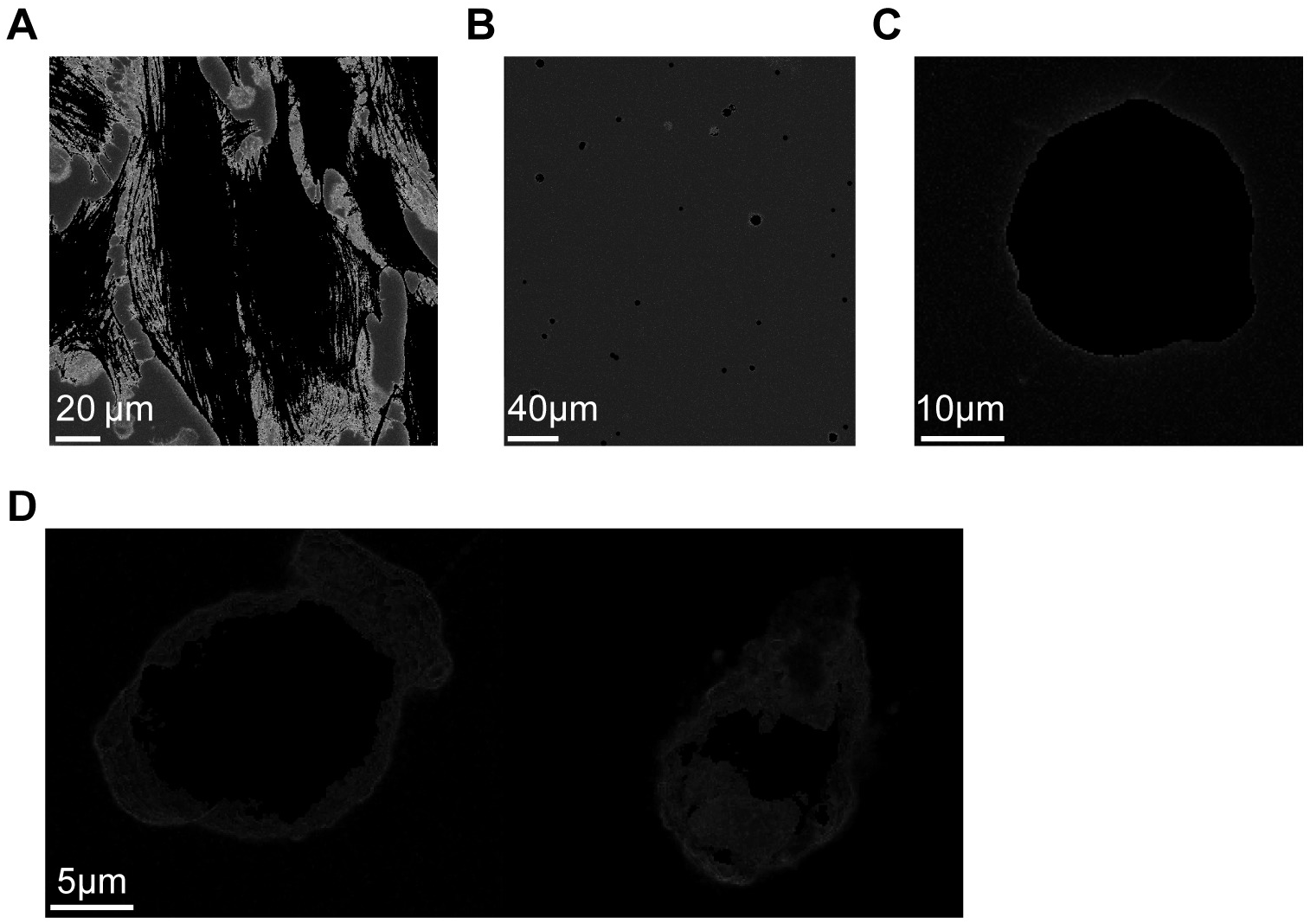
Separated background noise cluster images. (A) The separated background image by LUMoS of the BPAE cells image in Fig 3. (B) The separated background image by LUMoS of the multi-color beads image in Fig 4. (C) The separated background image by LUMoS of the colorful cell image in Fig 5. (D) The separated background image of the imaged cells in Fig 6.

**S7 Fig.**
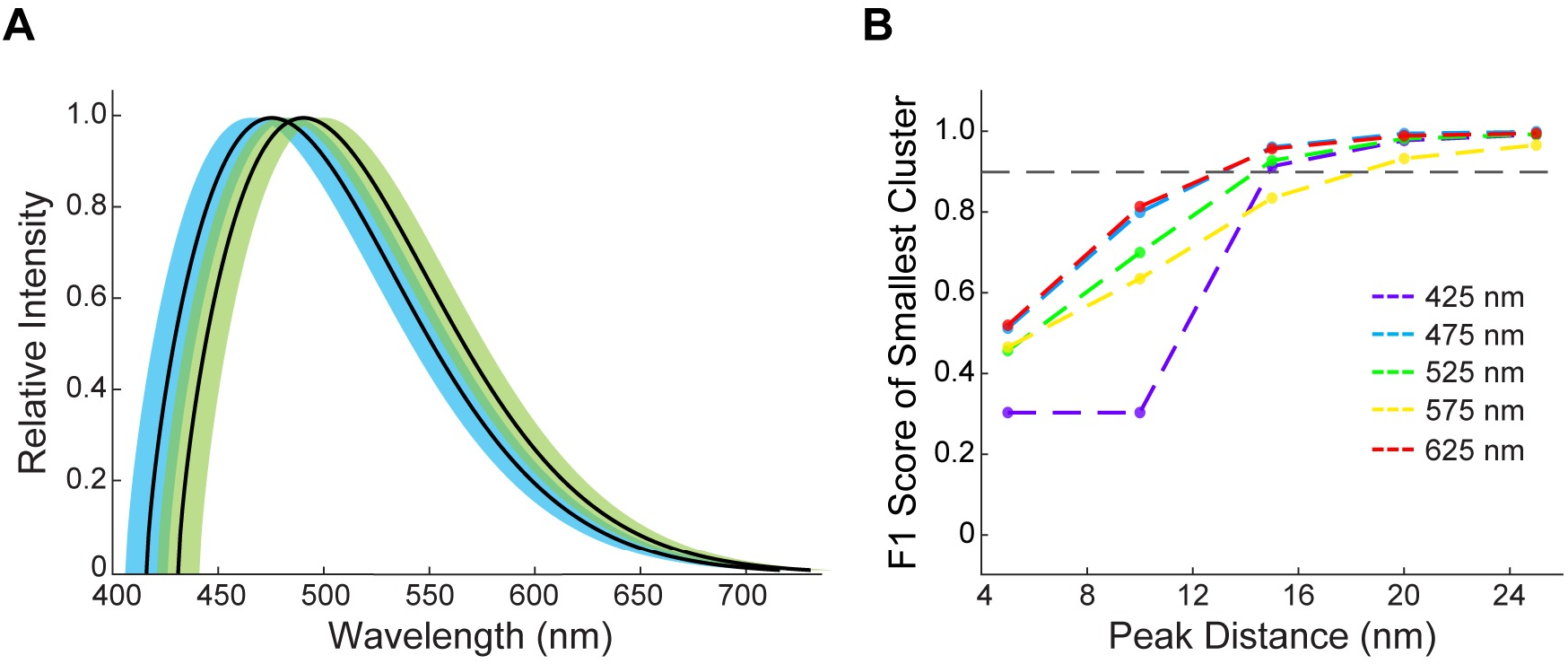
Two-fluorophore peak distance limitations. (A) Synthetic emission spectra of two fluorophores with peak emissions at 475 nm and 490 nm. 10 nm standard deviations for each spectra are shown in shaded area. (B) The performance of LUMoS for synthetic images of two fluorophores with variable distances between emission peaks. The cluster size ratio was fixed at 0.2, number of fluorophores at 2 and SNR at 10. For each color plotted, the peak of the lower wavelength fluorophore was fixed while the peak of the higher wavelength fluorophore was varied. Results of 10 simulations were averaged to obtain the final results.

